# MiR-101a loaded Extracellular Nanovesicles as Bioactive Carriers for Cardiac Repair

**DOI:** 10.1101/2020.02.28.970269

**Authors:** Jinli Wang, Christine J. Lee, Michael B. Deci, Natalie Jasiewicz, John M Canty, J. Nguyen

## Abstract

Myocardial infarction (MI) remains a major cause of mortality worldwide. Despite significant advances in MI treatment, many who survive the acute event are at high risk of chronic cardiac morbidity. Here we developed a cell-free therapeutic that capitalizes on the antifibrotic effects of micro(mi)RNA-101a and exploits the multi-faceted regenerative activity of mesenchymal stem cell (MSC) extracellular nanovesicles (eNVs). While the majority of MSC eNVs require local delivery via intramyocardial injection to exert therapeutic efficacy, we have developed MSC eNVs that can be administered in a minimally invasive manner, all while remaining therapeutically active. When loaded with miR-101a, MSC eNVs substantially decreased infarct size (12 ± 2.4% vs. 21.4 ± 5.7%) and increased ejection fraction (53.6 ± 7.6% vs. 40.3 ± 6.0%) and fractional shortening (23.6 ± 4.3% vs. 16.6 ± 3.0%) compared to control. These findings are significant as they represent an advance in the development of minimally invasive cardio-therapies.

## Background

Myocardial infarction (MI) remains a major cause of death in the US and beyond (1, 2). Even though advances in MI treatment by coronary reperfusion and/or systemic medication have significantly decreased mortality, many who survive the acute event continue to be at high risk of arrhythmias, congestive heart failure, and stroke (3). After MI, the myocardium undergoes extensive remodeling, with multiple processes affecting disease progression and healing of the damaged tissue.

Inflammatory monocytes and macrophages play important roles in disease progression and remodeling of the infarcted myocardium. Immediately after MI, macrophages infiltrate the infarcted myocardium, remove apoptotic cardiomyocytes, and degrade the extracellular matrix through the secretion of enzymes and cytokines (4). Subsequently, the injured tissue is replaced with non-contractile, fibrotic tissue and extracellular matrix, which increases myocardial stiffness and worsens cardiac function. Furthermore, fibrosis reduces blood flow through decreased capillary density and negatively affects mechanical strength (5). While established heart failure therapies including statins and aldosterone antagonists have minor anti-fibrotic effects, inhibiting fibrosis is not their main mode of action and anti-fibrotic mechanisms are poorly understood (6). Thus, no current therapy specifically and efficiently targets fibrotic tissue in the infarcted heart.

Transforming growth factor beta (TGF-β) signaling is extensively involved in cardiac fibrosis (7), so therapeutic agents capable of downregulating TGF-β would be highly desirable. MicroRNA (miR)-101a has been shown to target and modulate TGF-β and Wnt signaling, both of which are involved in fibroblast proliferation and activation and, therefore, fibrosis (8, 9). However, due to its anionic nature and large molecular weight, systemic delivery of naked miR-101a, as with other small RNAs, is ineffective and requires a delivery vehicle (10).

While macrophages are important for the removal of apoptotic cardiomyocytes in the initial phase post-MI, their excessive persistence can impair infarct healing. Aside from contributing to fibrosis, macrophages with a pro-inflammatory phenotype also secrete cytokines and chemokines that exacerbate myocardial inflammation (11). Since disease progression after MI is complex, a therapeutic system that simultaneously targets both cardiac remodeling and the inflammatory response after MI would be highly beneficial. Mesenchymal stem cell (MSC)-derived extracellular nanovesicles (eNVs) are one such promising therapeutic modality because they possess multifaceted regenerative effects. MSC eNVs possess intrinsic bioactivity and are anti-fibrotic, induce angiogenesis, and inhibit cardiomyocyte apoptosis (12, 13), all of which are pertinent to cardiac repair after MI.

The objective of this study was to exploit the antifibrotic effects of miR-101a and deliver it using a carrier with intrinsic tissue regenerative capacity, namely MSC eNVs (14). Carriers with intrinsic bioactivity are especially attractive in that they not only serve as drug vehicles but, unlike inert carriers, also exert therapeutic effects. One of the major drawbacks of current MSC eNV-based therapeutics is that they require direct intramyocardial injection to induce therapeutic effects. This is a highly invasive procedure that can lead to severe complications such as arrhythmia and tissue irritation. We showed that MSC eNVs mediate cardioprotective effects when they are loaded with anti-fibrotic miR-101a, even without direct myocardial injection. Intravenously administered miR-101-loaded MSC eNVs improve cardiac function in a mouse model of MI, have antifibrotic effects, and also polarize macrophages to an anti-inflammatory phenotype. These findings are significant as they represent an advance in the development of minimally invasive cardio-therapies.

## Methods

### Extracellular nanovesicle (eNVs) isolation

eNVs were isolated from human bone marrow-derived MSCs obtained from the American Type Culture Collection (ATCC PCS-500-012) and cultured with MesenPRO RS^™^ Medium (Gibco^™^, Gaithersburg, MD) (14-17). Passage number P1-P7 cells were used. After 2-3 days of MSC culture in EV-free medium, eNVs were isolated by differential centrifugation. Briefly, the cell culture medium was centrifuged at 2000 x g for 30 min to remove cells and debris. The supernatant was transferred to a new tube and mixed with EV isolation reagent and incubated overnight at 4°C. The eNV pellet was obtained by centrifugation at 10,000 x g for 1 h at 4°C and re-suspended in phosphate-buffered saline (PBS). eNVs were quantified by the Bradford assay at an absorbance of 595 nm.

### eNV characterization by transmission electron microscopy (TEM) and nanoparticle tracking analysis (NTA)

eNVs were prepared as previously described (14). Briefly, eNVs in PBS were air dried on type A carbon TEM grids (Ted Pella, Redding, CA) and stained with 1% uranyl acetate. After 5 min incubation, the remaining liquid was wicked away and the grids allowed to air dry. The grid was analyzed using a FEI Techni F20 G2 transmission electron microscope (FEI, Hillsboro, OR) at an accelerating voltage of 200 kV.

### Western blotting

eNVs were collected and isolated using Total Isolation Reagent (Thermo Fisher Scientific, Carlsbad, CA). The EV pellet was re-suspended and lysed with RIPA buffer and combined with 4× LDS buffer. Samples were heated to 95°C for 5 min, analyzed on a 4–12% Bis-Tris NuPAGE gel (Sigma Aldrich, St. Louis, MO), and transferred onto a PVDF membrane at 100 V for 60 min. Blots were incubated with anti-human CD63+ primary antibody (1:1000, Cat# 556019; Becton Dickinson, Franklin Lakes, NJ) or anti-human CD9+ primary antibody (1:1000, Cat# 555370; Becton Dickinson, Franklin Lakes, NJ) overnight at 4°C and visualized using the Bio-Rad ChemiDoc MP Imaging System (Bio-Rad Laboratories, Hercules, CA).

### Enrichment of miRNAs into eNVs

miRNA enrichment was performed using the Neon Transfection System (Thermo Fisher Scientific). Briefly, 240,000 MSCs were electroporated with 30 μg of the annealed miR-101a using the following settings: 990 V, 40 ms pulse, one pulse. MSCs were washed thoroughly with DMEM to remove free miRNA and cultured with EV-depleted medium. ENVs were collected 3 days after electroporation and analyzed for miR-101-3p content by RT-PCR using the qScript microRNA cDNA Synthesis Kit (Quanta Biosciences, Beverly, MA) and SYBR Mastermix (Quanta Biosciences) on a Stratagene Mx 3000 P (Stratagene, San Diego, CA). The primer for miR-101-3p was (TACAGTACTGTGATAACTGAA). A spike-in control was used for normalization.

### Fibrosis assay

Primary human cardiac fibroblasts were obtained from ScienCell (Cat# 6330, ScienCell Research Laboratories Inc., Carlsbad, CA). Passage number P3-P5 cells were used. Cells were plated onto a 24-well plate, cultured in fibroblast medium 2 (Cat# 2331, ScienCell) containing 5% FBS, and grown to 80% confluency. Cells were then serum starved for 12 h and treated with 0.5 - 2 μg of either unmodified or miR-101-enriched MSC eNVs. After 12 h, cells were stimulated with an additional 500 μl of complete medium containing TGF-β (Peprotech, Rocky Hill, NJ) at a final concentration of 10 ng/ml. To assess collagen production, we performed RNA extraction from the cells according to the TRIzol protocol (Thermo Fisher Scientific). cDNA was synthesized with the First Strand cDNA Synthesis Kit (E6300S; New England BioLabs, Ipswich, MA) and analyzed for expression of collagen I (primer sequences: FP GGGCAAGACAGTGATTGAATA and RP ACGTCGAAGCCGAATTCCT), and GAPDH (FP CAAGGTCATCCATGACAACTTTG and RP GTCCACCACCCTGTTGCTGTAG) by RT-PCR using the SYBR Mastermix (Quanta) on a CFX96 Touch^™^ Real-Time PCR Detection System.

### Bone marrow-derived macrophage isolation and culture

Bone marrow was harvested from femurs and tibias of C57BL/6 mice as previously described (18). After isolation and centrifugation, bone marrow cells were resuspended in DMEM/F12-10, frozen in DMEM/F12-40 with 10% DMSO and stored in liquid nitrogen until further use. Cells were thawed, washed once, and resuspended in macrophage complete medium (DMEM/F12, 10% FBS, 1% pen/strep, 100 U/ml recombinant murine M-CSF; Peprotech, Rocky Kill, NJ; Cat# 315-02). 5 ml of macrophage complete medium was added on day 3. On day 7, cells were used for polarization and immunofluorescence assays.

### Macrophage polarization assay

BMDM were treated with or without 6, or 16 μg/ml unmodified or miR-101 MSC modified eNVs. 24 h after treatment, pro-inflammatory and anti-inflammatory markers were quantified by RT-PCR (19). Briefly, cells were lysed with TRIzol reagent to extract RNA. cDNA was synthesized with the First Strand cDNA Synthesis Kit as above and analyzed for expression of iNOS (FP: CACCTTGGAGTTCACCCAGT, RP: ACCACTCGTACTTGGGATGC), IL-6 (FP: ACTTCACAAGTCGGAGGCTT, RP: TGGTCTTGGTCCTTAGCCAC), Arg1 (FP: GTGAAGAACCCACGGTCTGT, RP: CTGGTTGTCAGGGGAGTGTT), CD206 (FP: CAAGGAAGGTTGGCATTTGT, RP: CCAGGCATTGAAAGTGGAGT), and β-actin (FP: GCCTTCCTTCTTGGGTATGG, RP: CAGCTCAGTAACAGTCCGCC) by RT-PCR using iTaq SYBR Supermix (Bio-Rad, Cat# 1725125) on a CFX96 Touch^™^ Real-Time PCR Detection System (Bio-Rad). The same method was used for polarization assays on BMDMs. To assess the effects of exosomal surface proteins on macrophage polarization, BMDMs were treated with proteinase K pretreated MSC eNVs. Briefly, 10 μg/ml of proteinase K (PrtK) was used to treat MSC eNVs for 1.5 h at 37°C. Inactivation of PrtK was carried out by treatment with 0.1 mM phenylmethylsulfonyl fluoride (PMSF). Excessive proteinase K and PMSF were removed using molecular cutoff columns by three times buffer exchange using PBS. 16 μg/ml MSC ENVs were used to treat BMDM cells for 24 h followed by assessment of RT-PCR to measure M1/M2 related genes.

### BMDM immunofluorescence

BMDM cells were seeded into the cell imaging slide (Eppendorf, Cat# 0030742079) and incubated overnight at 37°C. Culture medium was replaced with fresh medium with or without 100 ng/ml LPS with 20 ng/ml IFN-g, 20 ng/ml IL-4, 16 μg/ml unmodified MSC eNVs, and 16 μg/ml miR-101 MSC eNVs.

After 48 h, samples were washed with PBS and fixed in 4% paraformaldehyde. Samples were then permeabilized with 0.25% Triton X-100 in PBS and blocked with 1% BSA PBS-T. Primary antibodies diluted 1:100 in 1% BSA were applied to the samples at RT for 1 h and washed. Samples were then incubated with secondary antibodies diluted 1:200 in 1% BSA at RT for 1 h. Cells were counterstained with DAPI and mounted with FluorSave mounting medium (Merck Millipore, Burlington, MA). The following antibodies were used: iNOS (pro-inflammatory marker; Abcam, Cambridge, UK; Cat#: Ab15323) and CD206 (anti-inflammatory marker; R&D Systems, Minneapolis, MN; Cat#: AF2535). Secondary antibodies were: AF568 (Invitrogen, Cat# A10042) and AF488 (Invitrogen, Cat# A11055).

### Animals

Male C57BL/6 mice were used (10-12 weeks) for the *in vivo* studies. All animal procedures were approved by the University at Buffalo - State University of New York Institutional Animal Care and Use Committee. All methods and experiments were performed in accordance with the U.S National Institutes of Health Guide for Care and Use of Laboratory Animals. Humane care and treatment of animals were ensured.

### Mouse model of myocardial infarction

MI was induced as previously described (19). Briefly, mice were anesthetized with isoflurane. Body temperature was maintained with a heating pad. A small incision under the mandible was made to visualize the trachea. Mice were intubated with a 20 gauge blunt needle and connected to a ventilator (Harvard Rodent Ventilator; Harvard Apparatus, Holliston, MA). A left lateral thoracotomy was made to expose the heart. The left anterior descending (LAD) artery was occluded permanently with an 8-0 nylon suture. The thorax was closed in layers (ribs, muscles, and skin). Analgesic treatments were provided according to the protocol and mice were monitored carefully.

### Biodistribution of MSC eNVs

For biodistribution studies (n = 3), 2 mg/kg of 1,1’-dioctadecyl-3,3,3’,3’-tetramethylindotricarbocyanine iodide (DiD)-labeled eNVs (n =3) or PBS (n=3) was administered via tail vein injection into C57BL/6 mice on day 2 after inducing MI. To fluorescently label the eNVs, DiD was dissolved in ethanol and incubated with eNVs at 37°C for 1 h. Ethanol and any unincorporated DiD were removed using the Pierce protein concentrator (3K) conditioned with PBS. Buffer exchanges were performed to remove unincorporated dye by centrifugation at 3000 x g at 4°C. After 5 h of injection, blood was collected by cardiac puncture under isoflurane anesthesia followed by centrifugation at 2000 rpm for 10 min to obtain the serum. Organs including brain, lung, heart, liver, spleen, kidneys, and bones were collected. EV biodistribution was analyzed using the IVIS Spectrum *in vivo* imaging system (PerkinElmer, Waltham, MA), and average region of interest (ROI) signals were calculated using Living Image 4.5.2 software (PerkinElmer).

### Immunohistochemistry - collagen staining

Collagen I was analyzed by immunofluorescence. Briefly, hearts were embedded in OCT, and cryosections (10 μm) were fixed with ice-cold acetone for 10 min. Samples were incubated with anticollagen I (Cat# ab34710, 1:100 dilution, Abcam) for 1 h at RT followed by Alexa Fluor 568-conjugated secondary antibodies (1:200 dilution, Invitrogen) for 45 min at RT. DAPI was used to counterstain nuclei. Sections at 90 μm intervals were analyzed. A total of 4 whole heart sections transversely from the apex to base at 900 μm intervals per heart were analyzed with n = 4 mice per group. Images were taken with a Zeiss AxioImager Microscope and analyzed using Image J.

### Immunofluorescence - staining for CD68, iNOS, and CD206

Ten μm frozen heart sections were fixed and permeabilized with ice-cold acetone for 10 min. After 3x washing with tris-buffered saline (TBS) containing 0.05% Tween-20 (TBST), tissue sections were blocked with protein block (Abcam) for 1 h at room temperature. Samples were incubated with the primary antibodies (1:100 dilution; CD68, Bio-Rad, Cat# MCA1957; iNOS, Abcam, Cat# ab15323; CD206, R&D Systems, Cat# AF2535) in TBST containing 2% BSA for 1 h at room temperature. Samples were washed in TBST 3x and incubated with Alexa Fluor 646-, Alexa Fluor 568-, or Alexa Fluor 488-conjugated secondary antibodies (Invitrogen; 1:200 dilution) for 45 min at RT. Samples were washed 3x with TBST, counter-stained with DAPI, washed 3x with TBST, mounted using FluorSave (Merck Millipore), and imaged using the Zeiss fluorescence microscope.

### Assessment of cardiac function and infarct size

To assess therapeutic efficacy, 2 mg/kg of unmodified and miR-101-loaded eNVs were administered to C57BL/6 mice via tail vein injection at day 2 and 3 after inducing MI (n=5 per group). Studies were performed in a blinded manner. Of the 15 animals randomized to treatment, 3 were excluded because of sudden death 2-3 days post MI [1 was PBS treated, 1 was treated with eNV, and 1 was treated with modified eNV). PBS-treated mice served as controls. Heart function was evaluated by echocardiography on day 11 after surgery. Mice were anesthetized by isoflurane (1-3%). Echocardiograms were performed with the GE Healthcare Echocardiography Vivid 7 system (GE Healthcare, Chicago, IL) equipped with an i13L probe. Two-dimensional mode parasternal long and short axis views were recorded. The left ventricular (LV) dimensions and wall thicknesses were determined from M-mode images at the mid-papillary muscle level. The LV stroke volume (SV) was calculated as the difference between the end diastolic volume (EDV) and the end systolic volume (ESV). LV ejection fraction (EF) was calculated as (EF=[(EDV-ESV)/EDV]*100). All measurements were performed with investigators blinded to experimental groups.

To determine infarct size, the same mice were sacrificed at day 13 after MI. Potassium chloride (30 mM) was injected into the LV to arrest the heart in diastole. Hearts were embedded in OCT and sliced transversely from the apex to base at 900 μm intervals. Heart sections (10 μm) were fixed with 10% neutral formalin solution and stained with the Masson Trichrome Stain Kit (Cat# 87019; Richard-Allan Scientific Co, San Diego, CA). Whole heart images were taken using the Zeiss AxioImager Microscope (Karl Zeiss). Infarct size was measured by Image J and calculated as: (1) Area measurement: infarct size = (sum of infarct areas from all sections)/(sum of left ventricle areas from all sections) x 100, (2) Midline length measurement: midline infarct length was defined as midline length of infarct that included more than 50% of the whole thickness of myocardial wall, and infarct size = (sum of midline infarct length from all sections)/(sum of midline circumferences from all sections) x 100 (20).

## Results

### Characterization of MSC eNVs

The hydrodynamic diameter of MSC eNVs measured by NTA and TEM was approximately 81 nm (**Figure 1A,B**), and eNVs were positive for CD63 and CD9 as expected (21) (**Figure 1C,D**).

**Figure 1.**
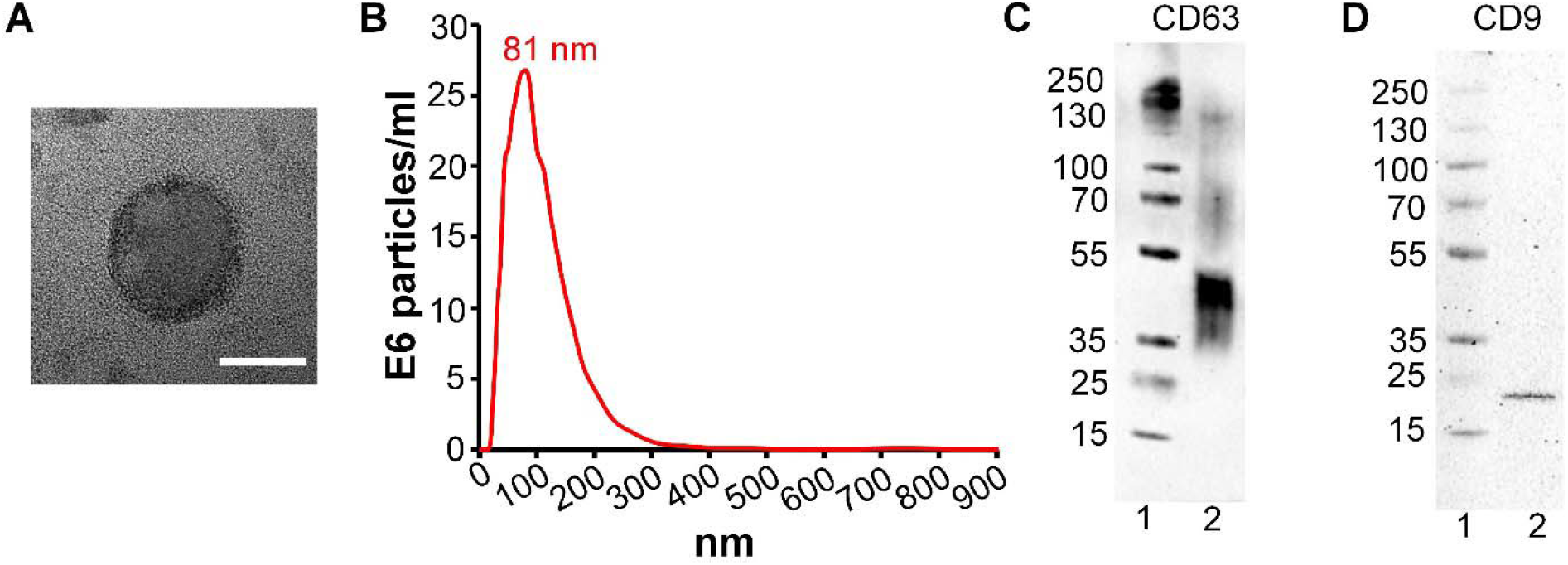
Characterization of MSC-derived eNVs. (**A**) TEM images of a representative MSC eNV, scale bar = 50 nm. (**B**) Size distribution of eNVs by NTA. (**C**) CD63 eNV surface marker protein expression by western blotting, protein ladder (lane 1) and eNV sample (lane 2) (**D**) CD9 MSC eNV surface marker expression by western blotting, protein ladder (lane 1) and eNV sample (lane 2).

### miR-101a loaded MSC eNVs inhibit collagen production

To further improve the intrinsic anti-fibrotic effects of MSC eNVs, MSC eNVs were loaded with the antifibrotic miR-101a (**Figure 2A,B**). miR-101a was selected because it has been shown to inhibit fibrosis through targeting TGF-β in ischemic cardiac diseases (8, 9). MSC eNVs electroporated with miR-101a using the virus-free electroporation approach achieved 315-fold enrichment compared to eNVs derived from unmodified MSCs (p < 0.05, **Figure 2B**). miR-101a-loaded MSC eNVs significantly downregulated collagen type 1A1 in TGF-β-stimulated cardiac fibroblasts compared to non-treated cells (**Figure 2C**).

**Figure 2.**
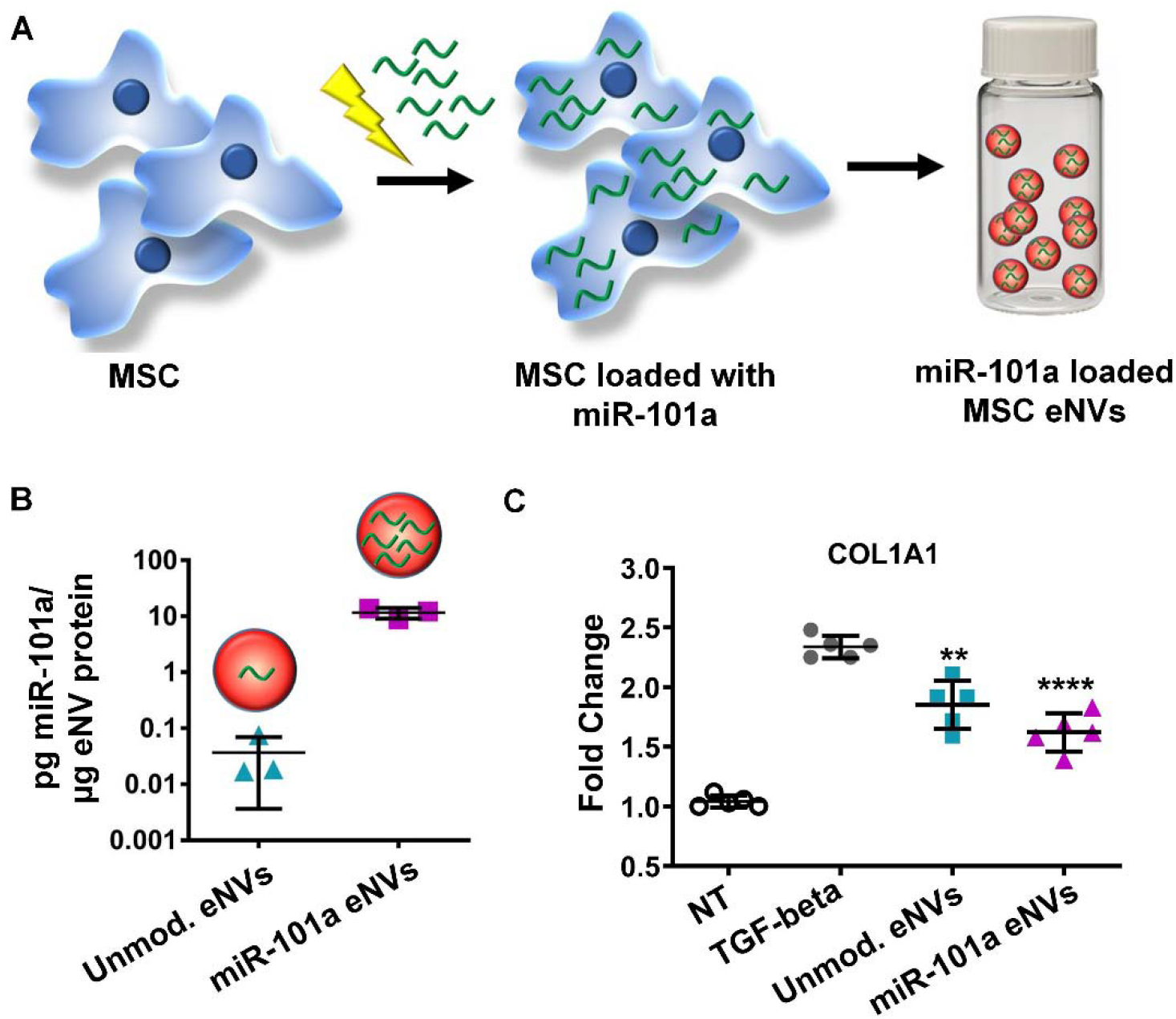
(**A**) eNVs derived from MSCs electroporated with miR-101 showed 315-fold enrichment with miR-101 than eNVs isolated from non-electroporated MSCs. (**B&C**) Collagen expression of cardiac fibroblasts treated with unmodified eNVs and miR-101 eNVs. Data are presented as mean ± SD with *p <0.05, **p<0.01, ****p<0.0001 by one-way ANOVA followed by Tukey’s post-test, n=4.

### Biodistribution of MSC eNVs and colocalization with CD68-positive macrophages *in vivo*

Next we assessed the biodistribution of DiD-labeled MSC eNVs after systemic administration via tail vein injection in a mouse model of MI. The majority of MSC eNVs were found in the spleen (64.8 ± 7.2%) and liver (56.7 ± 9.7%) followed by the lung (5.4 ± 1.3%), bone marrow (4.7 ± 1.3%), and kidneys (2.0 ± 0.7%) (**Figure 3A-C**). No brain deposition was detectable. About 3.8 ± 1% of MSC eNVs were found in the infarcted heart (**Figure 3B**), most likely as a result of the enhanced permeability of the leaky vasculature after MI (22). Importantly, MSC eNVs generally colocalized with macrophages present in the infarcted myocardium and spleen (**Figure 3 D,E**).

**Figure 3.**
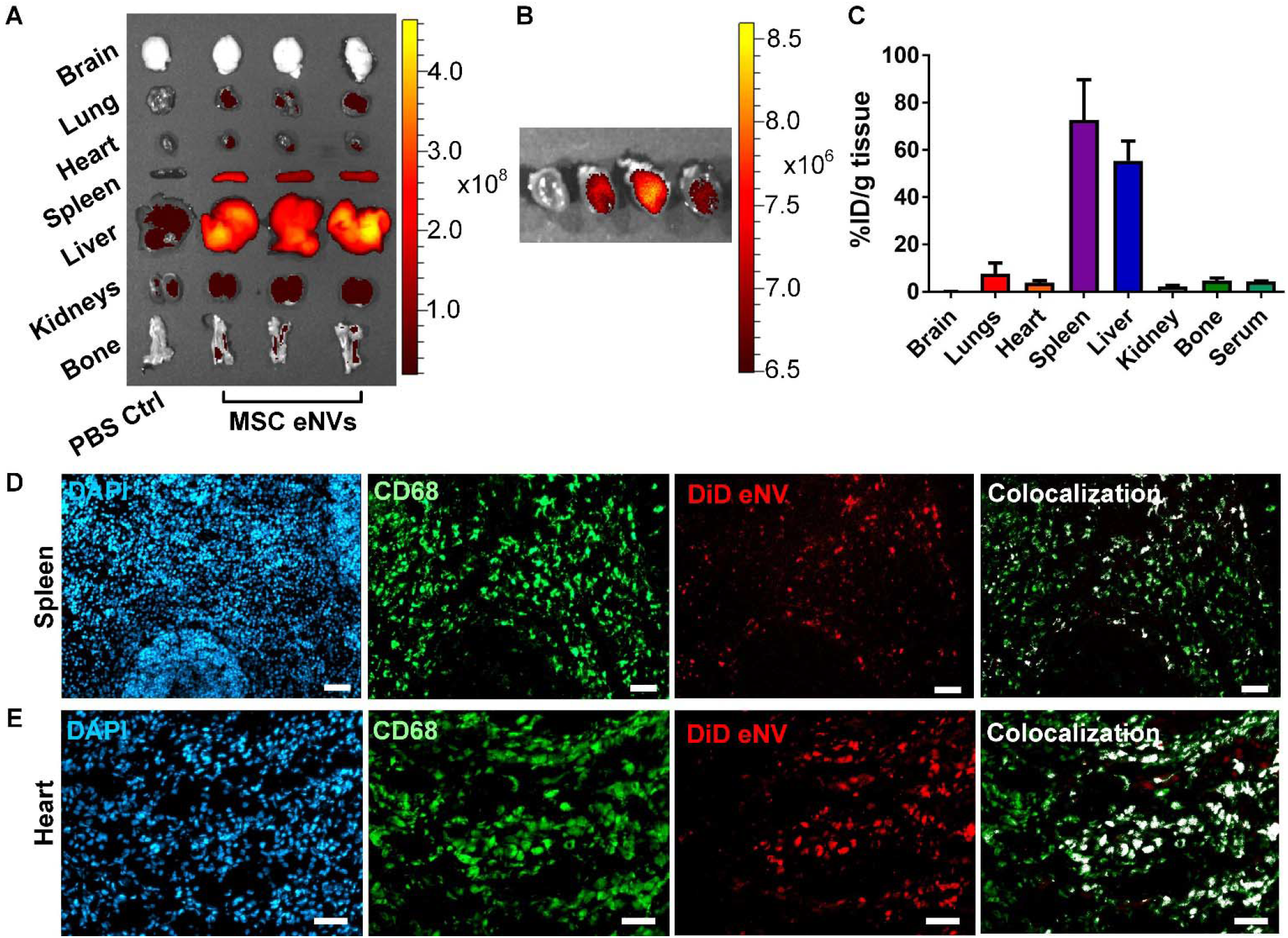
*In vivo* biodistribution of MSC-derived eNVs in mice with MI. (**A**) Representative images acquired using the IVIS Spectrum *in vivo* imaging system of DiD-labeled MSC eNVs in brain, lungs, heart, spleen, liver, kidneys, and bone. (**B**) IVIS imaging shows eNV deposition in the infarcted heart. (**C**) Quantification of % of full injection dose (% ID) in organs. Data are presented as mean ± SD, n=3. Colocalization studies from (**D**) splenic and (**E**) and cardiac tissues. Nuclei were stained with DAPI (blue), macrophages were stained with CD68 (green), eNV were stained with DiD (red), and colocalization of DiD-labeled eNV with CD68 are shown in white. Scale bar = 50 μm.

### MSC eNVs polarize macrophages to the anti-inflammatory phenotype

Because we discovered that MSC eNVs colocalized with macrophages, we examined the effects of MSC eNVs on macrophages. Depending on their microenvironment, macrophages can adopt different phenotypes, with the anti-inflammatory phenotype promoting wound healing and the inflammatory phenotype and contributing to disease progression (23). As shown in **Figure 4 A,B**, unmodified and miR-101 MSC eNVs increased the polarization of the bone marrow-derived macrophages to the antiinflammatory phenotype by 1.5 to 2-fold as measured by Arginase 1 and CD206 expression. No significant effects on pro-inflammatory macrophage markers were observed (iNOS and IL-6). In line with the gene expression markers quantified by RT-PCR, immunofluorescence staining showed that BMDMs treated with both unmodified MSC eNVs and miR-101a-loaded MSC eNVs were CD206 positive (**Figure 4C**). Similar to the cells treated with IL-4, a positive control for anti-inflammatory BMDMs, BMDMs treated with unmodified MSC eNVs and miR-101a-loaded MSC eNVs adopted a more spindle-like shape. By contrast, pro-inflammatory BMDMs stimulated with LPS and IFN-⍰ stained positive for iNOS and were significantly larger and more spherical in morphology compared to IL-4-stimulated BMDMs (**Figure 4C**).

**Figure 4.**
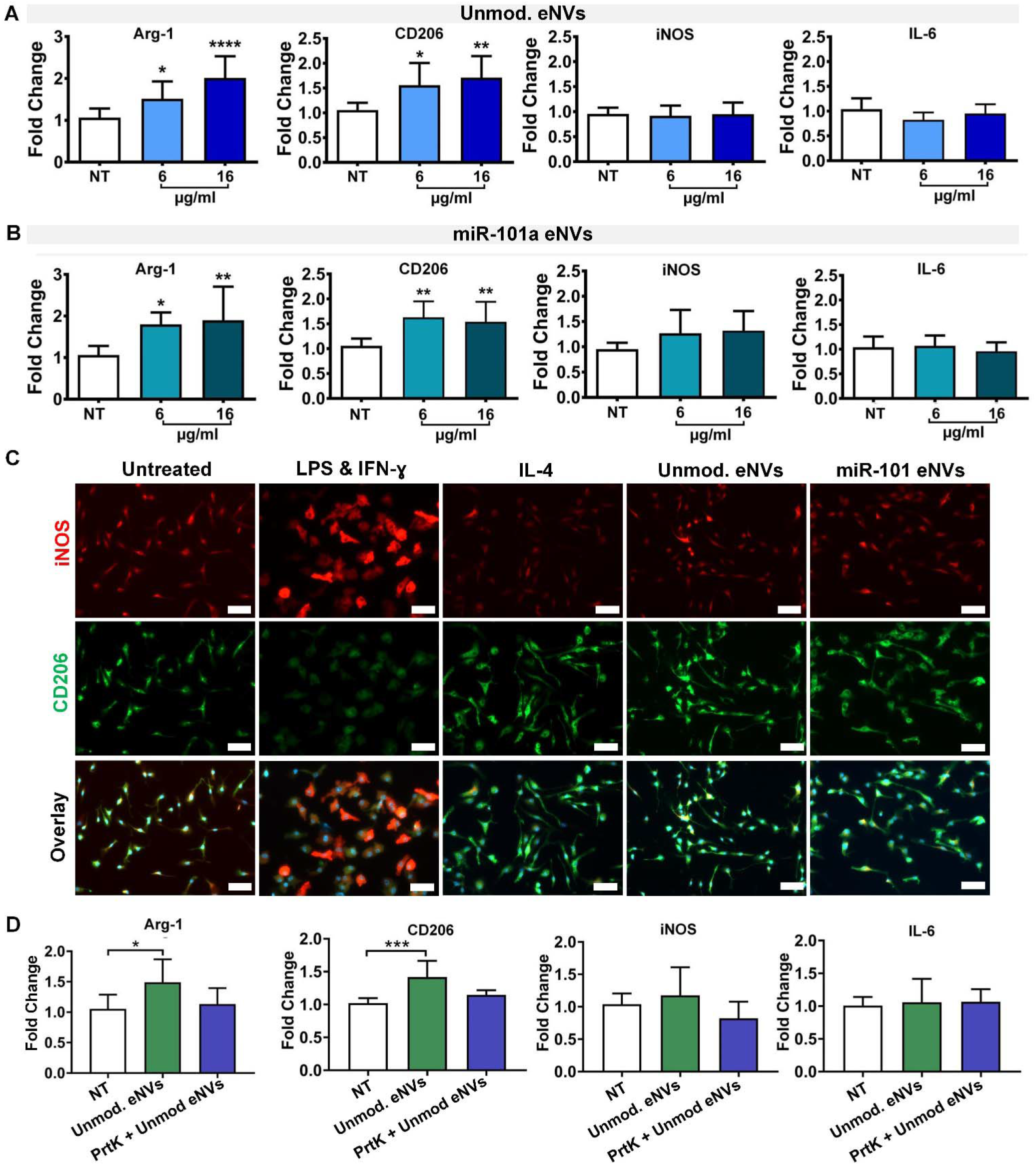
(**A, B**) Unmodified eNVs and miR-101 eNVs induce macrophage polarization to the antiinflammatory phenotype as measured by Arginase 1 and CD206 expression. No statistically significant differences in pro-inflammatory markers iNOS and IL-6 were observed. Data are presented as mean ± SD n=8 with *p<0.05, **p<0.01, and ****p<0.0001 with one-way ANOVA followed by Dunnett’s post-test. (**C**) Immunofluorescent staining of BMDMs treated with LPS and IFN-⍰, IL-4, unmodified MSC eNVs and miR-101 MSC eNVs. iNOS (red) served as a marker of the pro-inflammatory BMDM phenotype and CD206 served as a marker of the anti-inflammatory BMDM phenotype. Nuclei were stained with DAPI (blue). Scale bar: 50 μm. (**D**) Effect of proteinase K-treated BMDM eNVs on the gene expression of antiinflammatory markers Arg-1 and CD206 and pro-inflammatory markers iNOS and IL-6.

We next asked if proteins on the surface of the nanovesicles mediated some of the anti-inflammatory effects on BMDMs by treating MSC eNVs with proteinase K to digest all surface-bound proteins. Proteinase K-digested MSC eNVs did not upregulate anti-inflammatory-associated genes such as Arg-1 and CD206 (**Figure 4D**), indicating that surface proteins play a role in the anti-inflammatory effects of MSC eNVs.

### MSC eNVs increase the number of anti-inflammatory macrophages in the infarcted heart

After demonstrating that MSC eNVs polarized macrophages to the anti-inflammatory M2 phenotype, we determined the number of infiltrating CD68-positive monocytes and the percentage of antiinflammatory phenotype macrophages in the infarcted myocardium after treatment with MSC eNVs *in vivo.* To quantify the number of CD68-positive and anti-inflammatory macrophages, cardiac cryosections were stained with CD68 as a general macrophage marker, iNOS to identify pro-inflammatory macrophages, and CD206 to identify anti-inflammatory macrophages (**Figure 5**). Mice treated with unmodified MSC eNVs (56 ± 5% M2 macrophages) and miR-101-loaded MSC eNVs (53 ± 6% M2 macrophages) showed a significant increase in infiltrating anti-inflammatory macrophages in the infarcted area compared to mice treated with PBS alone (36 ± 2% M2 macrophages). With respect to the CD206/iNOS (M2/M1) ratio, PBS-treated mice had a significantly lower ratio (0.67 ± 0.1) in the infarcted area than mice treated with unmodified MSC eNVs (1.08 ± 0.26) and miR-101-loaded MSC eNVs (1.04 ± 0.09), indicating that unmodified MSC eNVs and miR-101-loaded eNVs have antiinflammatory effects. No differences were observed between the unmodified MSC eNVs and miR-101-loaded eNVs. This is not unexpected since the miR-101a is not known to target inflammatory processes.

**Figure 5.**
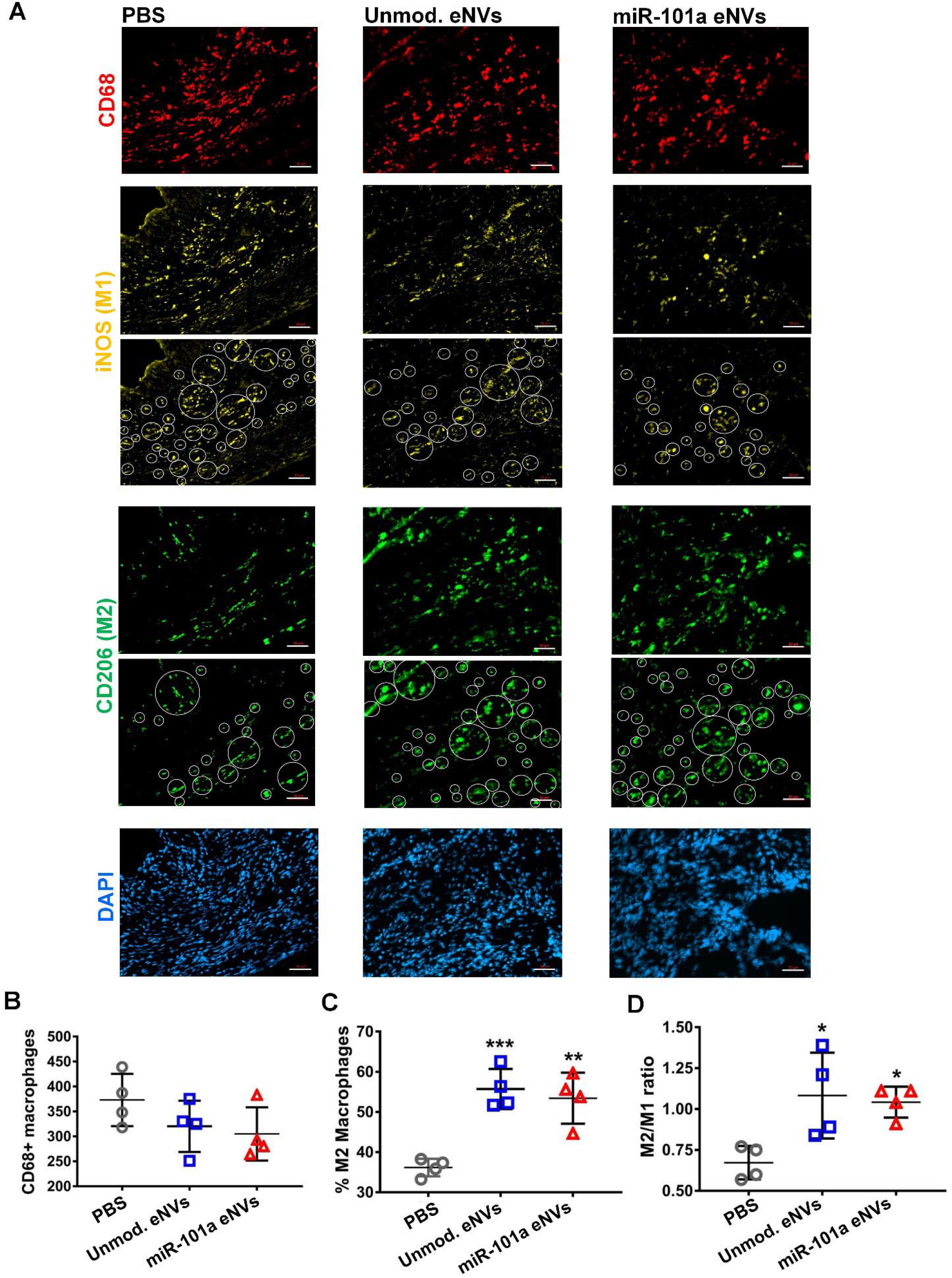
(**A**) Immunofluorescent staining of cryosections of the infarcted heart of mice treated with PBS, unmodified eNVs, and miR-101-loaded eNVs. Cryosections were stained for CD68 (red), the pro-inflammatory marker iNOS (yellow), the anti-inflammatory marker CD206 (green), and nuclei (DAPI, blue). White circles indicate iNOS and CD206 positive cells that were detected in the infarcted area by ImageJ. Quantification of (**B**) CD68-positive macrophages in the infarcted area, (**C**) % anti-inflammatory macrophages, and (**D**) M2/M1 (CD206/iNOS) ratio. Data are presented as mean ± SD (n = 4 per group). Images were taken from the infarcted areas.

### miR-101-loaded MSC eNVs decrease in vivo collagen production in the infarcted myocardium

To assess the in vivo effects of miR-101-loaded MSC eNVs and unmodified MSC eNVs on collagen production after MI, heart sections from mice treated with PBS, unmodified eNVs, and miR-101a-loaded MSC eNVs were stained with an anti-collagen antibody and assessed by immunofluorescence (**Figure 6A**). Representative images taken from the border zone, which was defined as the 2 mm area encircling the infarct area were shown (**Figure 6 A, B**). miR-101a-loaded MSC eNVs treated hearts had significantly less collagen (6.1 ± 0.3%) compared to mice treated with PBS (11.4 ± 1.5%; p<0.001) or unmodified eNVs (9.2 ± 1.0%; p<0.05) for the whole heart analysis (**Figure 6C**).

**Figure 6.**
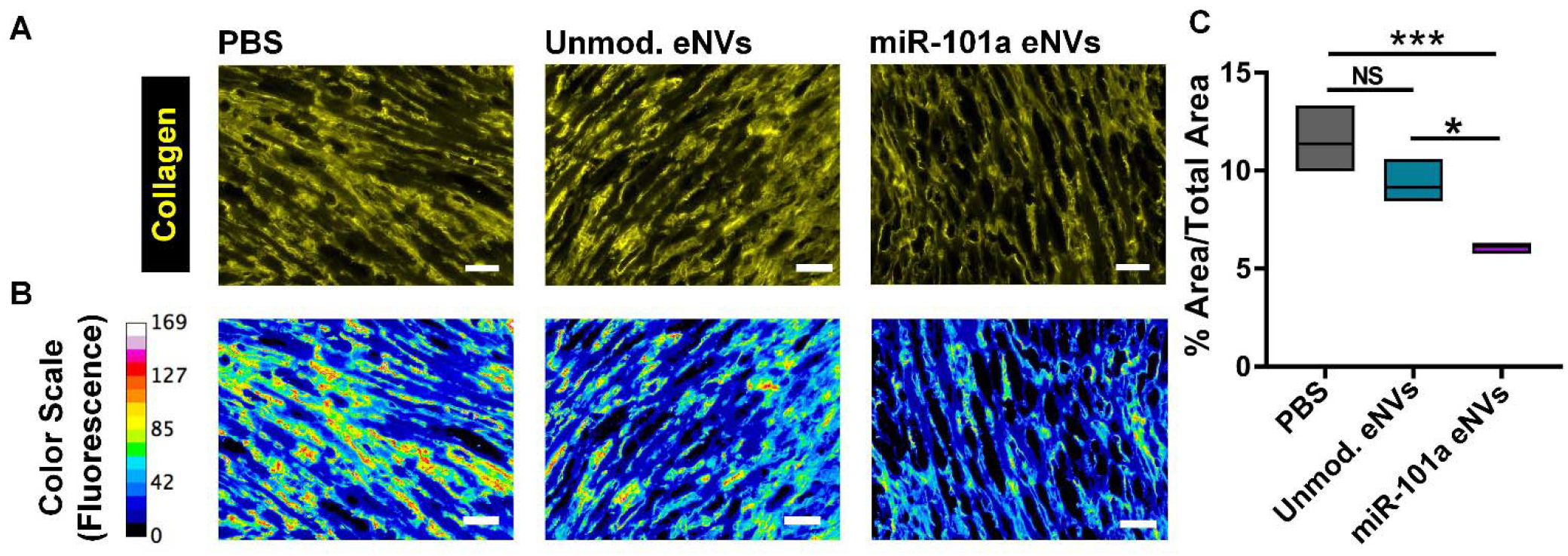
Effects of unmodified and miR-101 eNVs on collagen production. (**A**) C57BL/6 with induced MI were treated with unmodified eNVs or miR-101 eNVs and collagen I was stained with an anti-collagen I antibody with secondary visualization with Alexa Fluor 568 (yellow). Representative images from the border zones, which were defined as the 2 mm area encircling the infarcted area are shown. (**B**) To better visualize collagen staining intensity, a 16-color range function in ImageJ was used to indicate negative collagen staining (blue), yellow and orange indicate medium-intensity collagen staining, and red and pink indicate high-intensity collagen staining. (C) Quantification of collagen of the entire crosssectional area of the heart by Image J. Data are presented as mean ± SD (n =4) with *p<0.05, ***p<0.001 by one-way ANOVA followed by Tukey’s post-test.

### miR-101-loaded MSC eNVs display cardioprotection after acute MI

Finally, we assessed the effect of miR-101-enriched eNVs on myocardial function 13 days after MI. Unmodified eNVs showed a trend towards improving heart function by echocardiography and the improvements were not significant compared to PBS-treated mice (**Figure 7**). However, miR-101a-loaded MSC eNVs significantly decreased infarct size compared to PBS-treated controls analyzed by area-based (12 ± 2.4% vs. 21.4 ± 5.7%; p<0.05; **Figure 7B**) and length-based measurement (14.55 ± 2.66% vs. 31.89 ± 7.06%; p<0.01; **Figure 7B**). Mice treated with miR-101-loaded MSC eNVs displayed significantly higher EF (53.6 ± 7.6% vs. 40.3 ± 6.0% p<0.05; **Figure 7C**) and fractional shortening (23.6 ± 4.3% vs. 16.6 ± 3.0; p<0.05; **Figure 7C**) compared to mice treated with PBS. Other functional parameters assessed by echocardiography such as LVIDs and ESV (**Figure 7C**) were also significantly improved by miR-101-loaded MSC eNVs.

**Figure 7.**
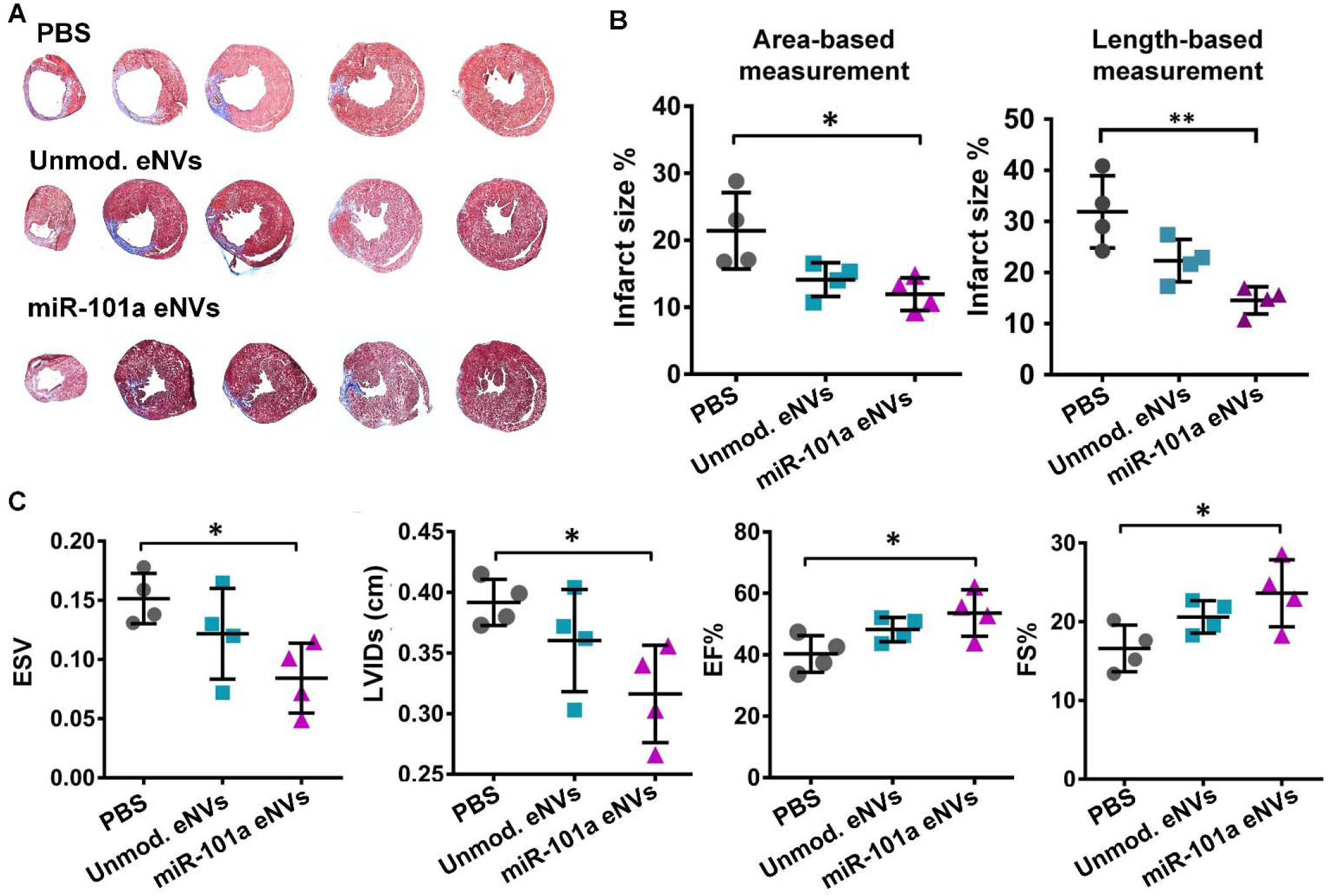
Heart function and infarct size after MI. (**A**) Masson trichrome staining of representative cryosections of infarcted hearts. (**B**) Infarct area expressed as % infarct size by area-based measurements (infarct area/LV (%)) and length-based measurements (midline infarct length/LV midline length (%)). (**C**) Analysis of ESV, LVIDs, ejection fraction (EF%), and fractional shortening (FS%) after MI by echocardiography. Data are presented as mean ± SD with *p < 0.05, **p < 0.01 by one-way ANOVA followed by Tukey’s post-test, n=4.

## Discussion

The current management of acute MI involves restoring cardiac perfusion through mechanical and drug interventions. Even when blood flow is quickly restored, the ischemic myocardium suffers injury and more efficient therapeutic approaches are required to fully restore cardiac function. In the first few days after MI, pro-inflammatory macrophages infiltrate the injured myocardium where they predominate as they clear cellular debris and degrade the extracellular matrix (24, 25). Pro-inflammatory macrophages also release TGF-β, which stimulates fibroblast proliferation and collagen secretion from cardiac fibroblasts (26). Although extracellular production of collagen is essential for cardiac repair, excessive collagen production leads to maladaptive cardiac remodeling. Timely reprogramming of inflammatory macrophages to an anti-inflammatory phenotype would, therefore, be important to myocardial healing.

While MSC eNVs have been tested for their cardio-regenerative therapy, the majority of studies use the invasive approach of direct intramyocardial injection of MSC eNVs to achieve high local concentrations at the affected site. For example, miR-132-loaded eNVs preserved heart function after direct transplantation into the heart (27). Similarly, miR-19a-enriched eNVs isolated from MSCs with GATA-4 overexpression reduced infarct size when injected to the ischemic border (28). Since direct injection into the myocardium is (a) highly invasive, (b) can lead to severe complications such as arrhythmia and tissue irritation, and (c) is often limited to a few injections, having a non-invasive route of administration could represent a significant advance in the development of next-generation cardio-therapies.

To develop a minimally invasive cardio-therapeutic, we loaded MSC eNVs with anti-fibrotic miR-101a to further improve on the reparative effects of MSC eNVs. miR-101a is a key inhibitor of fibrosis through targeting TGF-β. To overexpress miR-101a, studies have utilized viral transfection systems such as adenoviral transfection (8). However, the risk of immunogenicity with adenoviral gene therapies limits its broad clinical application, and indeed the inflammatory response evoked by adenoviruses can be fatal (29). MSC eNVs are highly attractive as a nucleic acid carrier for several reasons. First, MSC eNVs display low immunogenicity due to a lack of MHCII molecules and thus could be used as an allogeneic therapy. Second, the lipid bilayer of eNVs protects encapsulated nucleic acids against degrading enzymes. Third, unlike many synthetic carriers that are inert, MSC eNVs display intrinsic regenerative properties as they are anti-fibrotic, anti-apoptotic, and mediate angiogenesis (14).

We discovered, that enriching MSC eNVs with the anti-fibrotic miRNA miR-101 enhanced the therapeutic effect of MSC eNVs. Remarkably, treatment with miR-101-loaded eNVs significantly improved cardiac function and decreased infarct size compared to controls. The reparative effects of miR-101a loaded MSC eNVs compared to the unmodified eNVs appeared to be enhanced under *in vivo* conditions. Given that MSC eNVs have multifaceted effects, it is likely that additional, indirect mechanisms mediated by systemic exposure to MSC eNVs contribute to their overall function. Thus, we evaluated the effects of MSC eNVs on macrophage polarization. Pro-inflammatory macrophages are known to dominate in the early stage after MI and peak at 2-3 days after MI in humans and mice. A switch from the pro-inflammatory to the anti-inflammatory phenotype is required after MI to inhibit inflammation and promote wound healing (30). The persistence of pro-inflammatory macrophages has been shown to exacerbate heart injury due to persistent inflammation. When BMDMs were treated with unmodified eNVs and miR-101a loaded eNVs, anti-inflammatory markers such as CD206 and arginase-1 significantly increased whereas the effects on pro-inflammatory markers iNOS and IL-6 were negligible. Remarkably, mice treated with unmodified eNVs and miR-101a-loaded eNVs had increased numbers of anti-inflammatory macrophages (55% and 52%) compared to PBS treated control mice (38%). Because 50% of monocytes and macrophages dispatched to the infarcted myocardium originate from the spleen (31), it is likely that high splenic deposition after systemic administration of MSC eNVs is advantageous for modulating macrophage and monocyte function and contributes to the overall cardio-regenerative effect. Further supporting this, we found high colocalization of MSC eNVs with

CD68-positive macrophages, and Cho et al. demonstrated that MSCs have immuno-modulatory traits and accelerate cardiac repair by polarizing macrophages to the anti-inflammatory phenotype (32). MSC eNVs therefore share many features with their parent cells and are a promising alternative to cell-based therapeutics.

In conclusion, MSC eNVs serve as both a small RNA carrier and, by virtue of their intrinsic bioactivity, a therapeutic, making them a highly promising platform for cardiac therapy. Importantly, miR-101-loaded MSC eNVs significantly improved heart function, decreased infarct size, and decreased fibrosis after MI through anti-fibrotic and immunomodulatory effects. Notably, our studies showed that direct intramyocardial injection is not required for MSC eNVs to mediate cardio-protective effects when they are loaded with anti-fibrotic miR-101a. These results warrant further development of MSC eNV-based therapies for treating ischemic heart diseases.

## Acknowledgements

We would like to thank Dr. Joseph Spernyak (Roswell Park Cancer Institute) for technical assistance in the use of IVIS imager and funding by the NIH (S10 OD 016450). We thank Dr. Maixian Liu for the TEM images. Work in the Nguyen laboratory is supported by R01EB023262 (NIH), the Bruce Holm Technology award, and by the National Science Foundation (DMR 1751611). Work in the Canty laboratory is supported by HL-61610, the National Center for Advancing Translational Sciences UL1-TR-001412 and the Department of Veterans Affairs 1IO1BX002659.

## References

1. Finegold JA, Asaria P, Francis DP. Mortality from ischaemic heart disease by country, region, and age: statistics from World Health Organisation and United Nations. Int J Cardiol. 2013;168(2):934–45.

2. Naghavi M, Abajobir AA, Abbafati C, Abbas KM, Abd-Allah F, Abera SF, et al. Global, regional, and national age-sex specific mortality for 264 causes of death, 1980–2016: a systematic analysis for the Global Burden of Disease Study 2016. The Lancet. 2017;390(10100):1151–210.

3. Gerczuk PZ, Kloner RA. An update on cardioprotection: a review of the latest adjunctive therapies to limit myocardial infarction size in clinical trials. J Am Coll Cardiol. 2012;59(11):969–78.

4. Liehn EA, Postea O, Curaj A, Marx N. Repair After Myocardial Infarction, Between Fantasy and Reality: The Role of Chemokines. J Am Coll Cardiol. 2011;58(23):2357–62.

5. Travers Joshua G, Kamal Fadia A, Robbins J, Yutzey Katherine E, Blaxall Burns C. Cardiac Fibrosis. Circ Res. 2016;118(6):1021–40.

6. Fang L, Murphy AJ, Dart AM. A Clinical Perspective of Anti-Fibrotic Therapies for Cardiovascular Disease. Front Pharmacol. 2017;8:186–.

7. Dobaczewski M, Chen W, Frangogiannis NG. Transforming growth factor (TGF)-β signaling in cardiac remodeling. J Mol Cell Cardiol. 2011;51(4):600–6.

8. Pan Z, Sun X, Shan H, Wang N, Wang J, Ren J, et al. MicroRNA-101 Inhibited Postinfarct Cardiac Fibrosis and Improved Left Ventricular Compliance via the FBJ Osteosarcoma Oncogene/Transforming Growth Factor-β1 Pathway. Circulation. 2012;126(7):840–50.

9. Zhao X, Wang K, Liao Y, Zeng Q, Li Y, Hu F, et al. MicroRNA-101a Inhibits Cardiac Fibrosis Induced by Hypoxia via Targeting TGFβRI on Cardiac Fibroblasts. Cell Physiol Biochem. 2015;35(1):213–26.

10. Nguyen J, Szoka FC. Nucleic acid delivery: the missing pieces of the puzzle? Acc Chem Res. 2012;45(7):1153–62.

11. Frangogiannis NG. The inflammatory response in myocardial injury, repair, and remodelling. Nature reviews Cardiology. 2014;11(5):255–65.

12. Barile L, Lionetti V, Cervio E, Matteucci M, Gherghiceanu M, Popescu LM, et al. Extracellular vesicles from human cardiac progenitor cells inhibit cardiomyocyte apoptosis and improve cardiac function after myocardial infarction. Cardiovasc Res. 2014;103(4):530–41.

13. Lai RC, Arslan F, Lee MM, Sze NS, Choo A, Chen TS, et al. Exosome secreted by MSC reduces myocardial ischemia/reperfusion injury. Stem cell research. 2010;4(3):214–22.

14. Ferguson SW, Wang J, Lee CJ, Liu M, Neelamegham S, Canty JM, et al. The microRNA regulatory landscape of MSC-derived exosomes: a systems view. Sci Rep. 2018;8(1):1419.

15. Ferguson S, Kim S, Lee C, Deci M, Nguyen J. The Phenotypic Effects of Exosomes Secreted from Distinct Cellular Sources: a Comparative Study Based on miRNA Composition. AAPS J. 2018;20(4):67.

16. Ferguson SW, Megna JS, Nguyen J. 3 - Composition, Physicochemical and Biological Properties of Exosomes Secreted From Cancer Cells. In: Amiji M, Ramesh R, editors. Diagnostic and Therapeutic Applications of Exosomes in Cancer: Academic Press; 2018. p. 27–57.

17. Ferguson SW, Nguyen J. Exosomes as therapeutics: The implications of molecular composition and exosomal heterogeneity. J Control Release. 2016;228:179–90.

18. Amend SR, Valkenburg KC, Pienta KJ. Murine Hind Limb Long Bone Dissection and Bone Marrow Isolation. J Vis Exp. 2016(110).

19. Wang J, Seo MJ, Deci MB, Weil BR, Canty JM, Nguyen J. Effect of CCR2 inhibitor-loaded lipid micelles on inflammatory cell migration and cardiac function after myocardial infarction. INTERNATIONAL JOURNAL OF NANOMEDICINE. 2018;13:6441–51.

20. Takagawa J, Zhang Y, Wong ML, Sievers RE, Kapasi NK, Wang Y, et al. Myocardial infarct size measurement in the mouse chronic infarction model: comparison of area-and length-based approaches. J Appl Physiol (1985). 2007;102(6):2104–11.

21. Kowal J, Arras G, Colombo M, Jouve M, Morath JP, Primdal-Bengtson B, et al. Proteomic comparison defines novel markers to characterize heterogeneous populations of extracellular vesicle subtypes. Proceedings of the National Academy of Sciences. 2016;113(8):E968–E77.

22. Nguyen J, Sievers R, Motion JPM, Kivimae S, Fang QZ, Lee RJ. Delivery of Lipid Micelles into Infarcted Myocardium Using a Lipid-Linked Matrix Metalloproteinase Targeting Peptide. Mol Pharm. 2015;12(4):1150–7.

23. Hesketh M, Sahin KB, West ZE, Murray RZ. Macrophage Phenotypes Regulate Scar Formation and Chronic Wound Healing. Int J Mol Sci. 2017;18(7):1545.

24. Liu J, Wang H, Li J. Inflammation and Inflammatory Cells in Myocardial Infarction and Reperfusion Injury: A Double-Edged Sword. Clin Med Insights Cardiol. 2016;10:79–84.

25. Ong SB, Hernandez-Resendiz S, Crespo-Avilan GE, Mukhametshina RT, Kwek XY, Cabrera-Fuentes HA, et al. Inflammation following acute myocardial infarction: Multiple players, dynamic roles, and novel therapeutic opportunities. Pharmacol Ther. 2018;186:73–87.

26. Dean RG, Balding LC, Candido R, Burns WC, Cao Z, Twigg SM, et al. Connective tissue growth factor and cardiac fibrosis after myocardial infarction. J Histochem Cytochem. 2005;53(10):1245–56.

27. Ma T, Chen Y, Chen Y, Meng Q, Sun J, Shao L, et al. MicroRNA-132, Delivered by Mesenchymal Stem Cell-Derived Exosomes, Promote Angiogenesis in Myocardial Infarction. Stem Cells Int. 2018;2018:3290372.

28. Yu B, Kim HW, Gong M, Wang J, Millard RW, Wang Y, et al. Exosomes secreted from GATA-4 overexpressing mesenchymal stem cells serve as a reservoir of anti-apoptotic microRNAs for cardioprotection. Int J Cardiol. 2015;182:349–60.

29. Ritter T, Lehmann M, Volk H-D. Improvements in Gene Therapy. Biodrugs. 2002;16(1):3–10.

30. Nahrendorf M, Pittet MJ, Swirski FK. Monocytes: protagonists of infarct inflammation and repair after myocardial infarction. Circulation. 2010;121(22):2437–45.

31. Swirski FK, Nahrendorf M, Etzrodt M, Wildgruber M, Cortez-Retamozo V, Panizzi P, et al. Identification of Splenic Reservoir Monocytes and Their Deployment to Inflammatory Sites. Science (New York, NY). 2009;325(5940):612–6.

32. Cho DI, Kim MR, Jeong HY, Jeong HC, Jeong MH, Yoon SH, et al. Mesenchymal stem cells reciprocally regulate the M1/M2 balance in mouse bone marrow-derived macrophages. Exp Mol Med. 2014;46:e70.

